# HISAT-3N: a rapid and accurate three-nucleotide sequence aligner

**DOI:** 10.1101/2020.12.15.422906

**Authors:** Yun Zhang, Chanhee Park, Christopher Bennett, Micah Thornton, Daehwan Kim

## Abstract

Nucleotide conversion sequencing technologies such as bisulfite-seq and SLAM-seq are powerful tools to explore the intricacies of cellular processes. In this paper, we describe HISAT-3N (hierarchical indexing for spliced alignment of transcripts - 3 nucleotides), which rapidly and accurately aligns sequences consisting of nucleotide conversions by leveraging powerful hierarchical index and repeat index algorithms originally developed for the HISAT software. Tests on real and simulated data sets demonstrate that HISAT-3N is over 7 times faster, has greater alignment accuracy, and has smaller memory requirements than other modern systems. Taken together HISAT-3N is the ideal aligner for use with converted sequence technologies.

## Background

Nucleotide Conversion (NC) sequencing technologies are powerful tools that make chemical modifications to nucleic acids thereby gaining insight into highly dynamic cellular processes[1-3]. Bisulfite sequencing (BS-seq)[4] is one of the most well-known NC sequencing technologies. It uses bisulfite treatment to convert unmethylated cytosine to thymine in DNA molecules while leaving the methylated or hydroxymethylated cytosine (5mC or 5hmC) untouched (Table 1). BS-seq data has been used to qualitatively and quantitatively interrogate DNA methylation locations in genomic DNA for many years[4]. More recently, TET-assisted pyridine borane sequencing (TAPS)[5] technology has overcome the problem of producing low-complexity sequences which is present in the destructive bisulfite treatment of BS-seq. Another NC technique that has gained traction is known as thiol (SH)-linked alkylation for metabolic sequencing of RNA (SLAM-seq), which introduces 4-thiouridine (s^4^U) into living cells to replace uracil occasionally during the process of transcription in nascent RNA transcripts [6]. During the reverse transcription stage of the sample preparation process, the s^4^U is paired with guanine instead of an adenine, thus introducing guanine into the reverse strand. Sequencing reads are then generated with cytosine in place of thymine in the original sequence during PCR. Standard RNA-seq[7] can be used to detect adenosine-to-inosine RNA editing events by converting the modified adenosine (inosine) to guanosine during the reverse transcription. As sequencing technologies rapidly advance, NC technologies are likely to be combined with single cell sequencing technology such as scBS-seq[8] and scSLAM-seq[9], conveniently enabling researchers to understand various cellular processes at single-cell resolution. These NC sequencing technologies necessitate alignment strategies different from those commonly used aligners such as Bowtie2[10, 11], BWA[12], STAR[13], and HISAT2[14, 15], where converted nucleotides are treated as mismatches, often leading to the placement of sequences at incorrect genomic locations or failing to align them altogether. Inability to handle NC sequences leads to substantial bias, such as the omission of under-expressed genes prior to downstream analysis.

**Table 1.** Summary of several nucleotide conversion sequencing methods. BS-seq, TAPS, TAB-seq, and oxBS-seq are used to interrogate methylated, hydroxy methylated, or unmodified DNA cytosine. For example, BS-seq converts unmethylated cytosine to thymine while preserving methylated cytosine. SLAM-seq is used to interrogate nascent RNA transcripts, and RNA-seq can be used to interrogate adenosine-to-inosine RNA editing.

| Base | BS-seq | TAPS | TAB-seq | oxBS-seq | SLAM-seq | Standard RNA-seq |
| --- | --- | --- | --- | --- | --- | --- |
| C | T | C | T | T | C | - |
| 5mC | C | T | T | C | - | - |
| 5hmC | C | T | C | T | - | - |
| S<sup>4</sup>U | - | - | - | - | C | - |
| A | - | - | - | - | - | A |
| I | - | - | - | - | - | G |
| Base change | C -> T | C -> T | C -> T | C -> T | T -> C | A -> G |

To handle this erroneous alignment issue, several alignment programs were developed for aligning one specific type of NC sequencing data such as BS-Seq or SLAM-seq reads. Bismark[16], a widely adopted aligner primarily developed for BS-seq data, uses a three-nucleotide alignment strategy, i.e., the reference genome and sequencing reads are changed to use three-nucleotide prior to alignment. Bismark then uses Bowtie2 or HISAT2 as an underlying sequence aligner. These sequence aligners accurately handle reads that can be uniquely mapped, however they produce a randomly selected alignment when a read is mapped to multiple locations, with the mapping accuracy generally inversely proportional to the number of mappable locations. Since Bismark internally runs at least four instances of a sequence aligner (e.g., Bowtie2), it requires a great amount of memory (e.g., over 20 GB for human DNA sequences), and requires additional steps to handle and combine alignment output from the aligners, which is very time consuming. SLAM-DUNK[17], an aligner primarily developed for SLAM-seq reads, uses NextGenMap[18] as an underlying aligner. NextGenMap’s k-mer table-based alignment strategy enables fast mapping of reads, but NextGenMap is not designed to handle RNA-seq reads, especially those spanning multiple exons. This shortcoming of SLAM-DUNK could lead to incorrect alignment for such multi-exon spanning reads.

To date, no one has developed a unified methodology for aligning converted sequences and consolidating alignment of these technologies in one package. Each sequence aligner is tied to one specific technology with speed or accuracy problems. To overcome these limitations and improve speed and accuracy for NC read alignments, we have developed HISAT-3N, an extension of HISAT2 that implements a three-nucleotide alignment algorithm. HISAT-3N is the first general program that can handle any NC reads including BS-seq and SLAM-seq, RNA or DNA. HISAT-3N is substantially faster than other sequence aligners with higher mapping accuracy. Furthermore, HISAT-3N provides a tool to generate the conversion-table and identify the conversion status of nucleotides in read-level measurement. Users can study the dynamic cellular processes accurately and efficiently with HISAT-3N. Thus, HISAT-3N is the ideal option for handling three-nucleotide alignment. HISAT-3N is included in the HISAT2 package available at https://daehwankimlab.github.io/hisat2/hisat-3n.

## Results

We evaluated the performance of HISAT-3N and compared it to other commonly used NC aligners: Bismark and BS-seeker2[19] for BS-seq, and SLAM-DUNK for SLAM-seq. We used two data sets to evaluate HISAT-3N’s BS-seq alignment performance: (1) 10 million simulated paired-end DNA reads with a 50% C-to-T conversion rate and a 0.2% per-base sequencing error rate to evaluate the performance for non-directional whole genome bisulfite-seq read alignment, and (2) 78 million real paired-end whole genome BS-seq reads (SRA: SRR3469520[20]). Our test data sets for SLAM-seq alignment include (3) 10 million simulated single-end reads generated from 3’ regions of transcripts using realistic settings, e.g., with a 2% T-to-C conversion rate and a 0.2% per-base sequencing error rate and (4) 45 million real single-end SLAM-seq reads (SRA: SRR5806774[2]). The real reads are trimmed before alignment. We used the simulated data sets to estimate the alignment rate and accuracy of each aligner. We define alignment rate here as the number of aligned read pairs (or reads) divided by the total number of read pairs (or reads). Alignment accuracy refers to the number of correctly aligned read pairs (or reads) divided by the total number of read pairs (or reads). Multi-aligned read pairs (or reads) are classified as correctly aligned if any one of multiple alignments is correct.

HISAT-3N exhibits faster alignment speed and higher accuracy in simulated BS-seq data as compared to Bismark and BS-seeker2 (Table 2). HISAT-3N exhibits the fastest alignment speed, which is 7 and 23 times faster than Bismark and BS-seeker2, respectively, indicating that users could save considerable time by using HISAT-3N for mapping. In addition to speed, HISAT-3N also performs better in terms of alignment accuracy. Using only the 3N index, 98.52% of paired-end reads are mapped to the correct locations. Using both the 3N and repeat indexes improves alignment accuracy to 99.36% and requires only 10% more processing time. HISAT-3N also has high alignment accuracy for simulated SLAM-seq data (Table 3). SLAM-DUNK runs faster than HISAT-3N for alignment. However, the alignment rate and accuracy for HISAT-3N is substantially higher than for SLAM-DUNK. Although SLAM-DUNK has a similar alignment rate to HISAT-3N, 26.58% of the reads are partially mapped (not aligned end-to-end). Furthermore, because SLAM-DUNK cannot handle spliced alignment, there is a higher chance of aligning reads to incorrect positions. The alignment accuracy for HISAT-3N with repeat index reaches 99.81% versus 95.59% for SLAM-DUNK with 1,000 alignment searching.

**Table 2.** Performance comparison for HISAT-3N, Bismark, and BS-seeker2 on 10 million simulated 100-bp paired-end BS-seq reads.

| Sequence aligner | HISAT-3N | HISAT-3N<br>(repeat) | Bismark | BS-seeker2 |
| --- | --- | --- | --- | --- |
| Runtime | 13 minutes | 14 minutes | 97 minutes | 302 minutes |
| Alignment rate | 99.65% | 99.65% | 95.45% | 96.53% |
| Unique alignment rate | 95.29% | 95.30% | 95.45% | 96.53% |
| Alignment accuracy | 98.52% | 99.36% | 95.36% | 95.16% |
| Alignment accuracy (unique) | 96.20% | 96.22% | 95.36% | 95.16% |
| Memory usage | 9.5 GB | 10.6 GB | 16.5 GB | 16.7 GB |

**Table 3.**
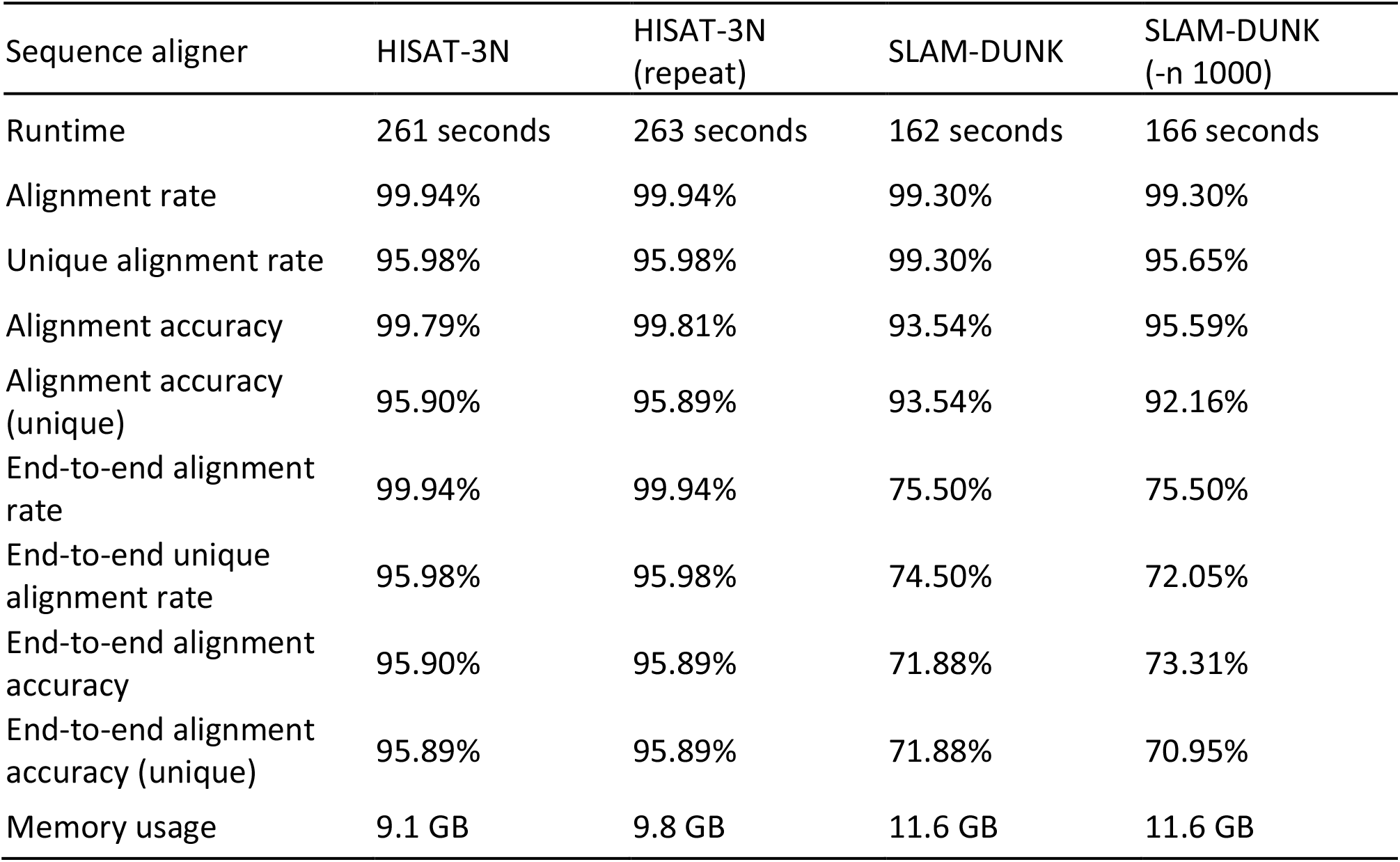
Performance comparison between HISAT-3N and SLAM-DUNK on 10 million simulated 100-bp single-end SLAM-seq reads. End-to-end alignment refers to mapping of 90% or more of a read’s sequence to the genome, or equivalently, at most 10% of a read’s sequence is soft clipped.

We also tested the alignment speed and rate for each sequence aligner on real data. When the input data size is large (for instance, 78 million paired-end reads), the speed advantage of HISAT-3N becomes clear (Table 4). HISAT-3N only took about 1.5 hours to map the data, while Bismark and BS-seeker2 required about 17 hours and 50 hours, respectively.

**Table 4.** Performance comparison for HISAT-3N, Bismark, and BS-seeker2 on 78 million real whole genome paired-end BS-seq reads.

| Sequence aligner | HISAT-3N | HISAT-3N<br>(repeat) | Bismark | BS-seeker2 |
| --- | --- | --- | --- | --- |
| Runtime | 87 minutes | 99 minutes | 1,031 minutes | 2,979 minutes |
| Alignment rate | 89.02% | 89.21% | 83.00% | 87.64% |
| Unique alignment rate | 85.19% | 84.77% | 83.00% | 87.64% |
| Memory usage | 8.4 GB | 10.2 GB | 16.5 GB | 16.7 GB |

When testing aligners with real SLAM-seq data (Table 5), we found that HISAT-3N maintained a high alignment rate (97.25%), though ran slightly slower than SLAM-DUNK. SLAM-DUNK aligned 99.17% of the reads when we set the maximum number of alignments to 1,000. Based on the results of our simulation reads, the alignment accuracy for SLAM-DUNK is only 95.59%, thus we infer that SLAM-DUNK results for the real reads could include many incorrect alignments.

**Table 5.** Performance comparison between HISAT-3N and SLAM-DUNK on 45 million real single-end SLAM-seq reads.

| Sequence aligner | HISAT-3N | HISAT-3N<br>(repeat) | SLAM-DUNK | SLAM-DUNK<br>(-n 1000) |
| --- | --- | --- | --- | --- |
| Runtime | 971 seconds | 1,055 seconds | 608 seconds | 987 seconds |
| Alignment rate | 97.25% | 97.25% | 99.17% | 99.17% |
| Unique alignment rate | 79.98% | 79.94% | 99.17% | 84.33% |
| Memory usage | 8.4 GB | 10.4 GB | 12.0 GB | 11.8 GB |

## Discussion

We have used and expanded HISAT2 index and alignment algorithms in order to develop HISAT-3N, which is specifically designed for the rapid and accurate alignment of NC sequencing reads. HISAT-3N has five key advantages over other NC aligners such as Bismark and SLAM-DUNK, making HISAT-3N versatile and easily employed: (1) HISAT-3N’s three-nucleotide alignment algorithm supports virtually any type of NC sample preparation protocol rather than being specific to one technology; (2) HISAT-3N uses the same index to map different types of NC reads (e.g., BS-seq, BS-RNA-seq, and SLAM-seq); (3) HISAT-3N seamlessly handles three-nucleotide DNA or RNA data; (4) HISAT-3N runs much faster due to bypassing the time intensive steps of writing and reading the intermediate alignment results from disk; and (5) HISAT-3N provides an option to report all multi-mapped reads regardless of the number of mapped places (e.g. 1,000 alignment positions), which considerably increases the alignment accuracy.

HISAT-3N uses the same conversion index for handling BS-seq, SLAM-seq, and any other conversion approaches as outlined in Table 1, thus making HISAT-3N independent of protocols and more versatile and convenient for use. To the best of our knowledge, all known NC sequencing technologies, including SLAM-seq, BS-seq, TAPS, TAB-seq[21], oxBS-seq[22], and standard RNA-seq, convert cytosine to thymine (or vice-versa) or convert adenine to guanine (or vice-versa), thus the REF-3N and REF-RC-3N indexes (Methods) are sufficient for unifying alignment of all known NC sequencing data types. HISAT-3N also allows for incorporation of splice sites, single-nucleotide polymorphisms, and small insertions/deletions using a graph index structure available in the HISAT2 program. Like most other sequence aligners, the HISAT-3N index only needs to be built once and is used for aligning samples of different kinds without the need to build other indexes.

The information reduction of the genome and reads that is characteristic of three-nucleotide algorithm obfuscates the true placement of the reads. Indeed, reads are mapped to more locations of the 3N reference compared to when the original reads are mapped to the original reference. We demonstrate this effect by aligning 10M 100-bp simulated single-end DNA reads. The four-nucleotide alignment with HISAT2 shows that only 5.65 % of reads can be mapped to more than one location. The three-nucleotide alignment with HISAT-3N shows 6.78% of reads are mapped to more than one location. Most aligners randomly report one or a subset of locations when reads are mapped to multiple locations. If users require an aligner to report all alignments, then the aligner may use 10 times more space for saving alignment outputs with a substantially increased runtime of the aligner. We apply HISAT2’s repeat index to resolve this multi-mapping problem (Figure 1). The whole genome index searching strategy (Figure 1A, left) needs to search each mapping position one by one, which can be very time consuming. The repeat index searching strategy (Figure 1A, right) can result in unique mapping, then expand the repeat mapping information to retrieve all the mapping positions. For multiple mapping, searching the repeat index can be substantially faster than the whole genome index. This repeat mapping process only adds 10% more runtime compared to when exclusively 3N indexes are used. More specifically, on a subset of simulated 10M single-end BS-seq reads, using the repeat index is three times faster than aligning reads using the two REF-3N indexes if the first 1,000 mapping locations are searched and outputted (Figure 1B).

**Figure 1.**
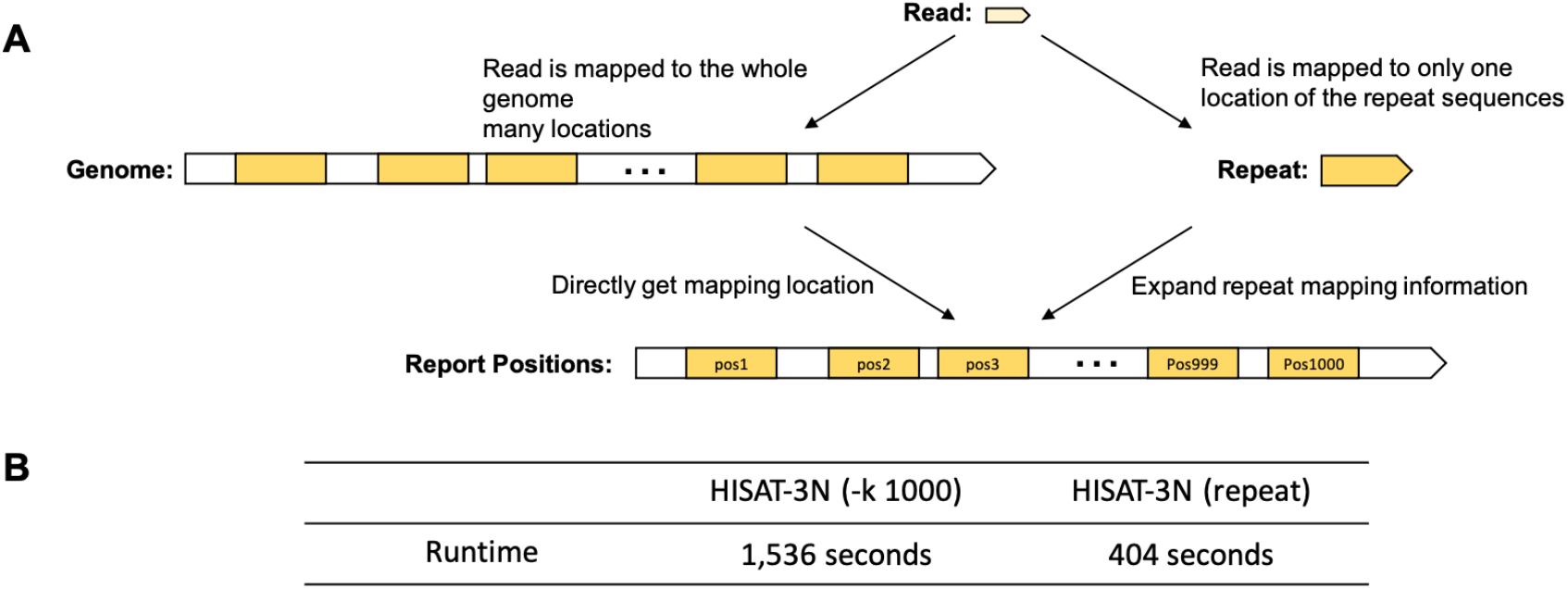
Repeat index enables faster three-nucleotide read alignment. HISAT-3N aligns reads using two different strategies: 1) HISAT-3N can directly align reads to the whole genome using the genome index and output their mapped locations (A, left) 2) HISAT-3N can use a repeat index to uniquely align reads to the repeat sequences regardless of how many locations to which they align on the genome (A, right). (B) Runtime comparison between direct mapping and repeat mapping strategy. The test data is 10M simulated single-end BS-seq reads (0.2% per-base sequencing error rate).

Not only is HISAT-3N between 7 and 23 times faster than other NC sequence aligners like Bismark and BS-seeker, HISAT-3N scales better than these aforementioned programs. When HISAT-3N multithreading is increased from 16 to 24 threads, HISAT-3N is 11- and 36-fold faster compared to Bismark and BS-seeker respectively (Table 6). Because Bismark and BS-seeker2 have to open four Bowtie2 simultaneously for the alignment process and only one thread for the alignment result filtration, the alignment speed is partially independent on the number of CPUs used. To use more CPUs for the filtering, Bismark and BS-seeker2 need to open four additional Bowtie2 and consume about 16 GB more memory, making them impractical for a common personal computer.

**Table 6.** Scalability comparison between HISAT-3N and SLAM-DUNK on 10 million simulated 100-bp paired-end BS-seq reads (0.2% per-base sequencing error rate).

| Sequence aligner | HISAT-3N | HISAT-3N (repeat) | Bismark | BS-seeker2 |
| --- | --- | --- | --- | --- |
| Runtime (16 threads) | 786 seconds | 842 seconds | 5,820 seconds | 18,142 seconds |
| Runtime (24 threads) | 514 seconds | 566 seconds | 5,689 seconds | 18,343 seconds |

## Conclusions

Here we present HISAT-3N, a rapid, versatile sequence aligner that processes reads generated by all known nucleotide conversion sequencing technologies including BS-seq, SLAM-seq, TAPS, oxBS-seq, TAB-seq, scBS-seq, and scSLAM-seq. HISAT-3N combines a three-nucleotide alignment strategy with the major alignment improvements present in HISAT2 to rapidly perform all nucleotide conversions in memory without storing and reading intermediate alignment results as is done by other aligners. This implementation substantially improves the processing speed and makes HISAT-3N an ideal choice for analyzing NC technologies’ data in the modern era.

## Methods

### Design principle

HISAT-3N’s three-nucleotide alignment method consists of 4 major steps: pre-alignment index building, read sequence conversion, three-nucleotide alignment, and result filtration. Here we use BS-seq data as an illustrative example without loss of generality, as other NC sequencing data types are similarly handled (Figure 2). HISAT-3N first builds two HISAT2 indexes from the reference human genome (GRCh38), denoted here as REF. The first index is built on REF with cytosine changed to thymine, denoted as REF-3N. The second HISAT2 index is built on the reverse complement of REF with cytosine changed to thymine, denoted as REF-RC-3N. While we have chosen cytosine-to-thymine to be our convention, the opposite conversion, i.e., thymine-to-cytosine, works equally well for handling BS-seq data.

**Figure 2.**
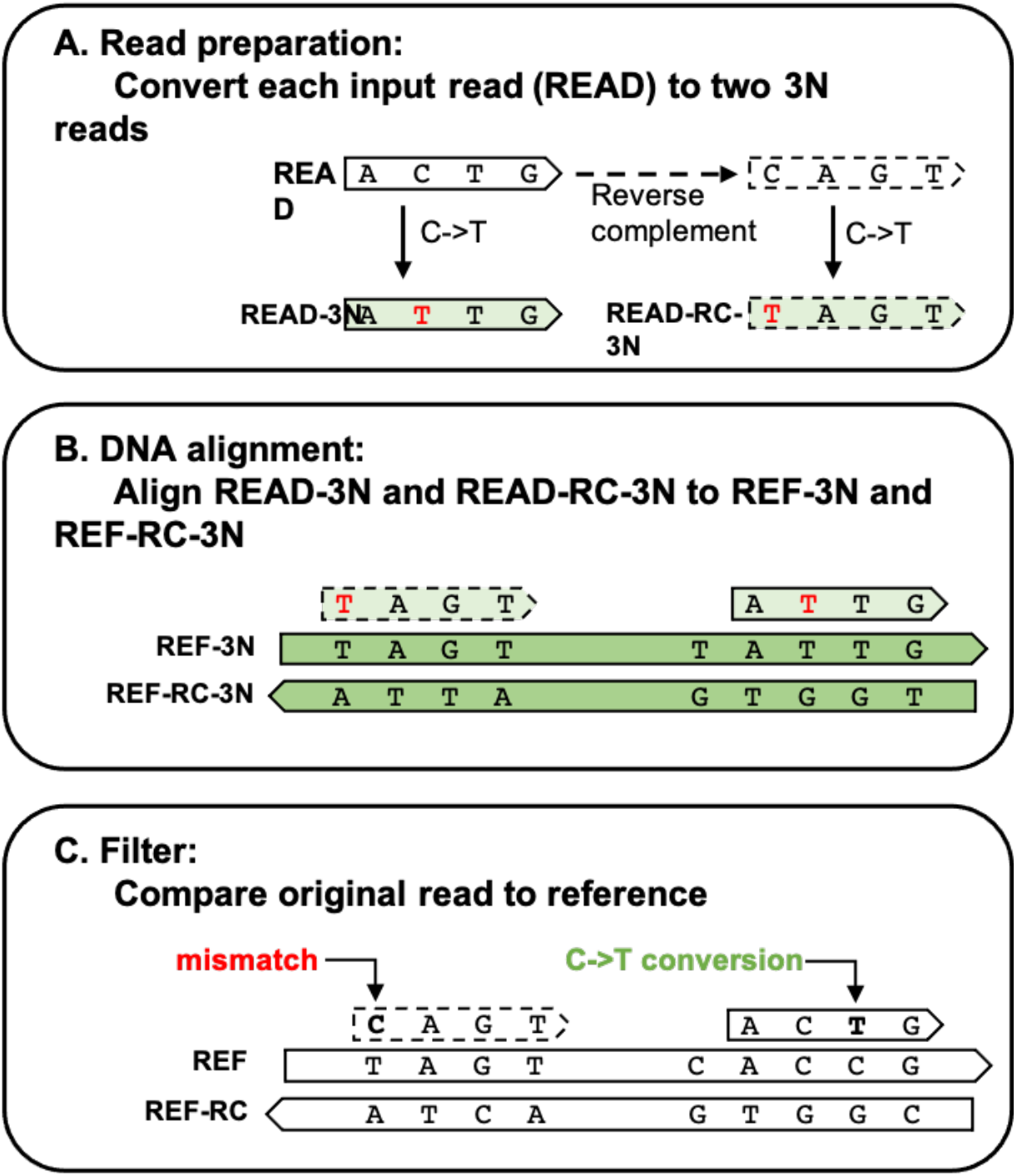
HISAT-3N alignment steps for BS-seq reads. (A) HISAT-3N converts each input read (READ) to two 3N reads: READ-3N and READ-RC-3N. READ-3N is READ with all thymine replaced by cytosine. READ-RC-3N is the reverse complement of READ, plus the replacement of cytosine with thymine. (B) HISAT-3N maps the two 3N reads to both REF-3N and REF-RC-3N references using pre-built indexes. (C) After the three-nucleotides alignment, HISAT-3N compares the original read sequence (READ) to the original four-nucleotides references (REF and REF-RC) to identify unmethylated cytosine positions and re-calculate an alignment score accordingly.

HISAT-3N uses FASTA or FASTQ formatted sequencing reads as inputs that can be compressed or uncompressed, single-end or paired-end. For each read, HISAT-3N generates two 3N reads (Figure 2A): (1) READ-3N: read with cytosine changed to thymine, and (2) READ-RC-3N: read that is first converted to its reverse complement, followed by changing cytosine to thymine. Note that in general READ-3N and READ-RC-3N are not reverse complements of each other.

Then HISAT-3N uses HISAT2 to align two versions of a read, READ-3N and READ-RC-3N, to both REF-3N and REF-RC-3N, involving a total of four alignment processes (READ-3N to REF-3N, READ-3N to REF-RC-3N, READ-RC-3N to REF-3N, and READ-RC-3N to REF-RC-3N) (Figure 2B). Depending on sequencing data types, HISAT-3N performs non-spliced alignment for DNA-seq reads (e.g., BS-seq) and spliced alignment for RNA-seq reads (e.g., SLAM-seq) (Figure 2B and Supplementary Figure 1, respectively).

After HISAT-3N aligns 3N reads to the 3N references, HISAT-3N compares the original read, READ, to the original genome reference, REF, for each alignment of 3N reads in order to identify converted nucleotide locations and mismatches (Figure 2C). HISAT-3N then sorts the alignment results by alignment score (a function of the number of mismatches and indels) and reports the alignments with the best alignment score in the SAM format[23]. During the output process, HISAT-3N adds extra SAM tags (Supplementary Note) to indicate the number of conversions and the aligned strand (REF or REF-RC). HISAT-3N directly performs this alignment readjustment of nucleotide conversions and mismatches directly in computer memory without storing and reading intermediate alignment results. Thus, this post-processing step of HISAT-3N is relatively fast, taking less than 10% of its total runtime. HISAT-3N also provides a script to create a 3N-conversion-table for methylated and unmethylated cytosines drawing from the SAM alignment file, as described in Supplementary Note.

### Repeat index and alignment

To build a repeat index, a set of identical sequences that are >= 40 bp and found in at least 5 locations of REF-3N and REF-RC-3N are combined into one repeat sequence. During the alignment process, if a read or its partial segments are mapped to 5 or more locations using 3N indexes (Figure 1), the read will be directly mapped to the repeat sequences using the repeat index, resulting in one repeat alignment per read. Then, HISAT-3N expands the repeat alignment result and outputs the alignment information in standard SAM format. The repeat index and HISAT-3N’s alignment algorithm enables reporting of all alignments for reads originating from repetitive genomic regions.

### Testing environment

We tested the alignment programs using a computer system with 24 CPU cores (two Intel Xeon E5-2680 v3) and 256 GB of memory. Each aligner is configured to use 16 threads so that each thread is assigned one CPU core on this system. Bismark and BS-seeker2 run four instances of Bowtie2 for the alignment process, and each instance of Bowtie2 uses 4 threads, resulting in a total of 16 threads.

### Reads adapter trimming

We trimmed the adapter sequence on real reads using trim-galore[24] (https://github.com/FelixKrueger/TrimGalore) and filtered out reads shorter than 40 bp to generate our real BS-seq and SLAM-seq data. We also trimmed poly-A sequences of the real SLAM-seq reads using trim-galore (Supplementary Note).

## Acknowledgements

This work was supported in part by the National Institute of General Medical Sciences (NIH) under grants R01-GM135341 and by the Cancer Prevention Research Institute of Texas (CPRIT) under grant RR170068 to D.K. All authors read and approved the final manuscript.

## Authors’ contributions

Y.Z., C.P., C.B., M.T., and D.K. performed the analysis and discussed the results of HISAT-3N. Y.Z., C.P., and D.K. designed and implemented HISAT-3N. Y.Z. and C.P. performed the evaluations of the various programs. Y.Z., C.P., C.B., M.T., and D.K. wrote the manuscript.

## Competing interests

The authors declare no competing financial interests.

## Availability of data and materials

Project name: HISAT-3N

Project home page: https://daehwankimlab.github.io/hisat2/hisat-3n

Operating system(s): Linux and Mac OS X

Programming language: C++, Python, Perl, and Java

License: GPLv3 license

## Notes

### Competing Interest Statement

The authors have declared no competing interest.

### Summary of Updates

Correct the home page address for HISAT-3N.

