## Supplemental Methods for "HISAT-3N: a rapid and accurate three-nucleotide sequence aligner"

### HISAT-3N: a rapid and accurate three-nucleotides sequence aligner
#### Supplementary Note

**Supplementary information:**

1. **Extra SAM tags**

**Supplementary method:**

1. **Software version**
2. **Command used to generate index**
3. **Command used to test BS-seq reads**
4. **Command used to test SLAM-seq reads**
5. **Command to generate 3N-conversion-table**
6. **Command to trim real reads**

#### Supplementary information:

1. **Extra SAM tags:**

   Yf:i:<N>
   Number of conversions are detected in the read.

   YZ:A:<A>
   The value + and – indicate the read is mapped to REF-3N (+) or REF-RC-3N (-).

#### Supplemental Method:

1. **Software version:**
   SLAM-DUNK version: 0.3.4
   Bismark version: 0.22.2
   BS-Seeker2 version: 2.1.8
   Bowtie2 version 2.3.4.3
   Trim-Galore (Cutadapt): 0.6.4

2. **Command used to generate index:**

   HISAT-3N index building:
   ```
   ./hisat-3n-build --3N -repeat-index 100-300 `reference
   genome fasta` `output tag`
   ```

   Bismark index building:
   ```
   ./bismark_genome_preparation --path_to_aligner `bowtie2
   directory` --bowtie2 `reference genome directory`
   ```

   BS-Seeker2 index building:
   ```
   python bs_seeker2-build.py -f `reference genome fasta` --
   aligner=bowtie2 `bowtie2 directory`
   ```

3. **Command used to test BS-seq reads (Table 2 and Table 4).**

   **HISAT-3N:**
   ```
   hisat-3n --index `HISAT-3N index` -p 16 --base-change C,T -
   -no-spliced-alignment -f -1 `BS-seq read 1` -2 `BS-seq read
   2` -S `output name`
   ```

   -f is for fasta format reads. To align fastq format, use -q rather than -f.

   **HISAT-3N (repeat):**

```
hisat-3n --index `HISAT-3N index` -p 16 --base-change C,T –
repeat --no-spliced-alignment -f -1 `BS-seq read 1` -2 `BS-
seq read 2` -S `output name`
```

**Bismark:**

```
bismark -f --path_to_bowtie2 `bowtie2 directory` --
non_directional --sam --bowtie2 -p 4 -genome `reference
genome directory` -1 `BS-seq read 1` -2 `BS-seq read 2`
```

**BS-Seeker2:**

```
python bs_seeker2-align.py -1 `BS-seq read 1` -2 `BS-seq
read 2` -g `reference genome` --aligner=bowtie2 `bowtie2
directory` -d `index directory` -f sam --bt2-p 4 -t Y
```

#### 4. Command used to test SLAM-seq reads (Table 3 and Table 5).

**HISAT-3N:**

```
hisat-3n --index `HISAT-3N index` -p 16 --base-change T,C -
f -U `SLAM-seq read` -S `output name`
```

-f is for FASTA format reads. To align FASTQ format, use -q rather than -f.

**HISAT-3N (repeat):**

```
hisat-3n --index `HISAT-3N index` -p 16 --base-change T,C -
-repeat -f -U `SLAM-seq read` -S `output name`
```

**SLAM-DUNK:**

```
slamdunk map -r `genome reference fasta` -o . -t 16 -5 0
`SLAM-seq read`
```

**SLAM-DUNK (-n 1000):**

```
slamdunk map -r `genome reference fasta` -o . -t 16 -5 0 -n
1000 `SLAM-seq read`
```

#### 5. Command to generate 3N-conversion-table

```
hisat-3n-table -p `n threads` --sam `sam file` --ref
`genome reference fasta` --table-name `outout name` --base-
change C,T
```

#### 6. Command to trim reads

**Trim adaptor:**

```
trim-galore -a --length 40 -o `output directory` `read
file`
```

**Trim polyA:**

```
trim-galore -a A{10} --length 40 -o `output directory`
`read file`
```
