## Supplemental figure for "HISAT-3N: a rapid and accurate three-nucleotide sequence aligner"

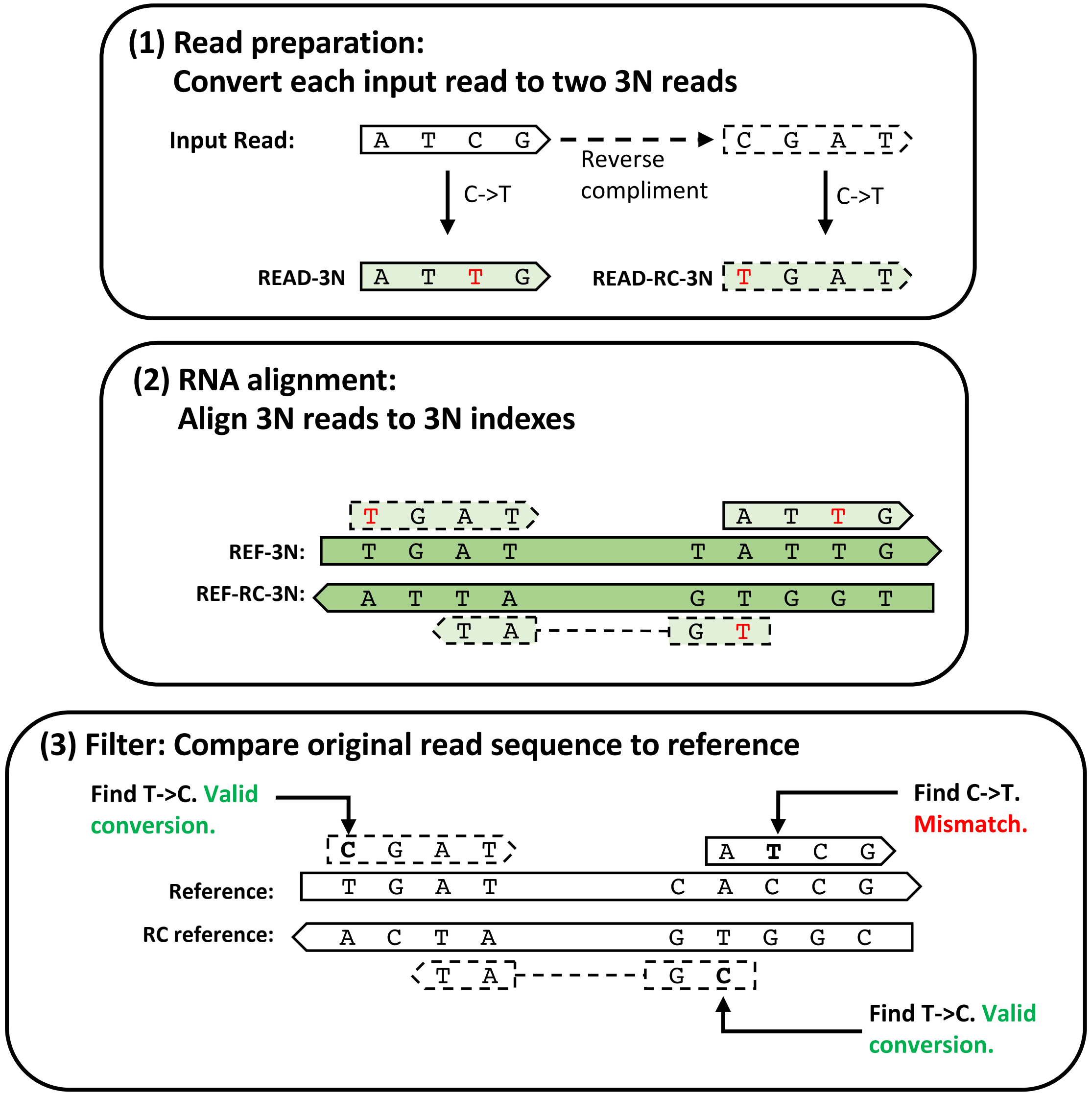


Supplementary Figure 1. Analysis workflow in HISAT-3N for SLAM-seq reads.
(A) HISAT-3N converts each input read (READ) to two 3N reads: READ-3N and READ-RC-3N. READ-3N is READ with all thymine replaced by cytosine. READ-RC-3N is the reverse complement of READ, plus the replacement of cytosine with thymine. (B) HISAT-3N maps the two 3N reads to both REF-3N and REF-RC-3N references using pre-built indexes. (C) After the three-nucleotide alignment, HISAT-3N compares the original read sequence (READ) to the original four-nucleotide references (REF and REF-RC) to identify unmethylated cytosine positions and re-calculate an alignment score accordingly.
